# The global dispersal of visceral leishmaniasis occurred within human history

**DOI:** 10.1101/2024.10.30.621037

**Authors:** João Luís Reis-Cunha, Cooper Alastair Grace, Sophia Ahmed, Simon E. Harnqvist, Cián M. Lynch, Mariana Côrtes Boité, Gabrielle Barcellos, Laurence Lachaud, Patrick Bastien, Harry Munt, Jeremy C. Mottram, Elisa Cupolillo, Daniel C. Jeffares

## Abstract

The *Leishmania donovani* species complex (LdSC) causes visceral leishmaniasis (VL), is present in Africa, Europe, the Middle East, Asia and the Americas, and causes 20-40,000 fatalities per year. Previous analyses concluded that dispersal of this species complex occurred 1-10 million years ago. Using updated methods and data, we show that a ten-thousand year old East African population dispersed globally only within the last 2,000 years, consistent with human migration, war and colonisation as driving factors.

## Main text

Genomic evidence has transformed our understanding of the history of human pathogens^1–3^, but relatively little is known about the evolutionary history of the protozoans that cause visceral leishmaniasis (VL), *L. donovani* and *L. infantum*. While the primary disease burden is in East Africa, the Indian subcontinent and Brazil, this species complex is now widely distributed, exposing 1.7 billion people to risk of infection and resulting in 20-40,000 fatalities/year ^4,5^.

Here we examine when the dispersal of these parasites occurred, using population genetic analyses based on genomic sequences from 899 isolates of the *Leishmania donovani* species complex from 35 countries, enabling the detection of 496,704 segregating single nucleotide polymorphisms (SNPs) (~15 Segregating sites/Kb). Unsupervised strain clustering indicated that the LdSC can be divided into 9 populations, with three main clades: East Africa, South America and South Asia; with intermediate/admixed groups in the Middle East, North Africa and Europe that confirm the admixture of *L. donovani* and *L. infantum* as an interbreeding species complex^6^ (Figure 1, Supplementary Figure 1, Supplementary Figure 2, Supplementary Figure 3, Supplementary Table 1).

**Figure 1:**
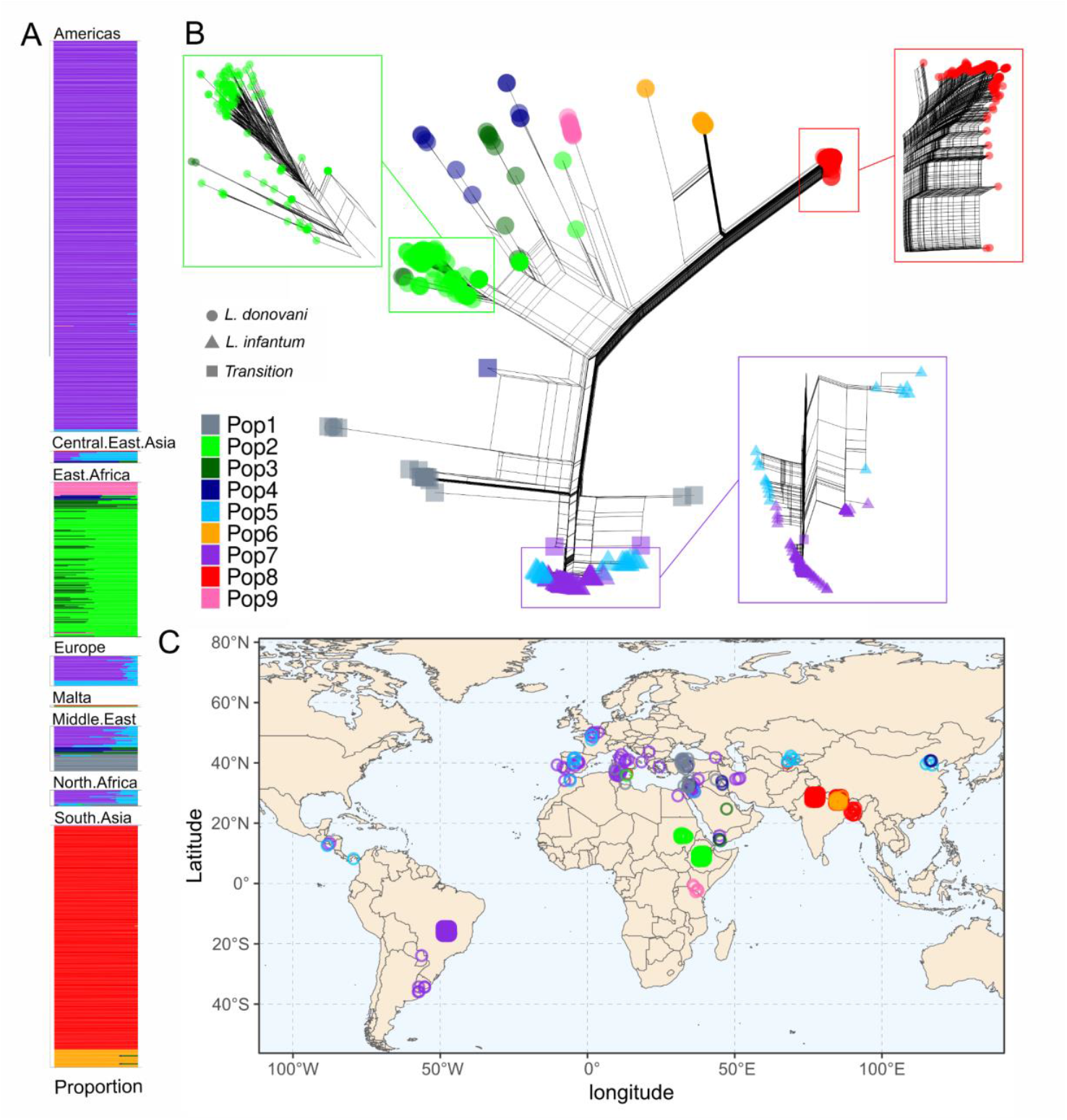
LdSC population structure reveals three main populations and intermediate isolates. Phylogeny and population structure based on the nuclear genome sequences of LdSC isolates. **A)** Admixture plot of the evaluated isolates, using K=9 (other K values can be seen in the Supplementary Figure 2). Each line corresponds to an isolate, and the proportion of each population is represented by colours. **B)** Phylogenetic network based on the nuclear genome sequences of LdSC isolates, coloured based on their main Admixture population assignment (as in **A**). Densely packed branches are zoomed and expanded to ease visualisation. *L. donovani, L. infantum* and *L. donovani/L. infantum* transition isolates are respectively represented by circles, triangles and squares. **C)** Map representing the country of isolation of each LdSC sample, coloured by its main Admixture population. The proportion of each population in each zone can be seen in Supplementary Figure 3).

While this and previous descriptive analyses of LdSC populations ^7–11^ portray genetic similarities within and between populations, they do not provide explicit timescales for the dispersal of the LdSC. To investigate this demographic history, we combined estimation of population coalescent times using mitochondrial DNA data, referred to as kinetoplast (kDNA) in trypanosomatids, accounting for population size variation^12^; with the estimates of population size variation and population split dates using nuclear SNP data produced with the sequential markov coalescent implemented in SMC++ ^13^. These analyses used two genomic regions (kDNA and nuclear), with different methods.

BEAST coalescent dating using kinetoplastid DNA SNPs from 567 of the 899 total isolates with a known date of isolation (17,024 nucleotides with 382 polymorphic sites, which are sites that are polymorphic in the population, ~22.4 segregating sites/Kb, mutation rate ~2.1×10^−6^ substitutions per nucleotide per year) indicated that the time to the most recent common ancestor (TMRCA) of all evaluated LdSC modern strains was in East Africa, at 355 CE (95% highest posterior density interval, HPDI 155 BCE to 836 CE) (Figure 2 A and B, Supplementary Figure 4). This is consistent with previous analysis implicating Sudan/Ethiopia as the earliest endemic region ^14–16^; with the root position in kDNA maximum likelihood phylogeny using closely related *Leishmania* species as outgroups (Supplementary Figure 4); and with the high genetic diversity of LdSC in Africa when compared with other populations (Figure 1A)^11^. This estimated period of LdSC dispersion from East Africa to other regions, coincides with the Aksumite rule in today’s Eritrea and Ethiopia during the first seven centuries CE ^17,18^. The Aksumites had extensive trade links with the Roman empire in the Mediterranean, the Iranian (Sasanian) empire, the Indian subcontinent and important links across the Red Sea into South Arabia (roughly today’s Yemen). These include trade and cultural connections but also periods of military intervention and invasion, such as the Aksumite invasion of the Himyarite kingdom in South Arabia in the 520s CE. These intercontinental trade links and conflicts could have facilitated movement of LdSC parasites, as migration of refugees and war increases VL incidence in modern times ^19,20^. Coalescent analysis of population split dates from kDNA indicated that the LdSC dispersed gradually to the Middle East, India and Europe over the following two thousand years, with the second major endemic area in South Asia arising in ~872 CE (95% HPDI 450 CE to 1280 CE) (Figure 2 A and B). *L. infantum* is now present in southern Europe, and is extremely abundant in Brazil.

**Figure 2:**
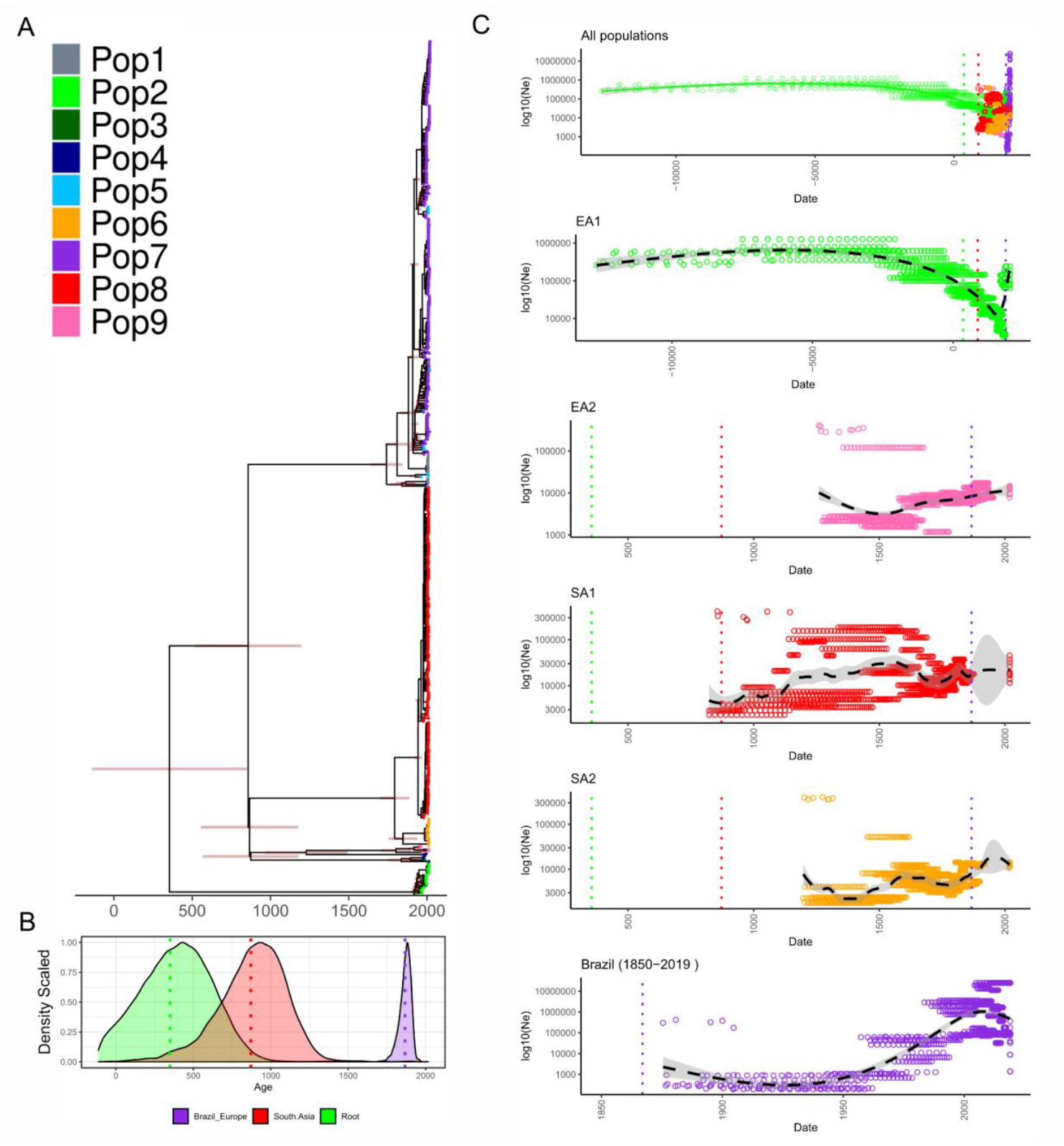
LdSC dating and population size through time. **A)** kDNA dated phylogeny of the LdSC isolates, where each isolated was coloured by their nuclear genome Admixture main population. **B)** Density plot representing the MCMC samplings for the full set (green curve - Root); South Asia coalescence point (red curve, Bayesian skyline coalescent prior, 3 population size estimates, GTR, fixed clock, 100M MCMC replicates); and the Brazil-Europe coalescence date (purple curve, exponential growth coalescent prior, GTR, fixed clock, 100M MCMC replicates). The green, red and purple dotted line represents respectively the BEAST mean root date for “Tree root” (355 CE; Africa), South Asia (872 CE) and Brazil-Europe (1867 CE). **C)** LdSC SMC++ population size estimate through time, using nuclear genome SNP data. The number of generations was converted to years, using the rate of 33.6 generations/year (see methods). Each panel corresponds to a different population. The green, red and purple dotted lines correspond to the same dates as in “B”. The oldest date for each population: EA1 12,680 BCE; EA2 1,247 CE; SA1 822 CE; SA2 1196 CE; Brazil 1873 CE; was supported by at least five from the 10 SMC++ replicates runs, with independent sampling. NE: Effective population size.

Previous analysis indicated that the *L. infantum* population in Brazil was derived from Spain, Portugal or France ^8^, consistent with our admixture analysis showing the genetic similarity between South America and European strains (Figure 1). BEAST estimation of coalescent time between the South American and Europe populations indicates that the sampled Brazilian lineages of *L. infantum* arose around 1867 CE (95% HPDI 1648 CE - 1941 CE), well after the initial contact of Europeans with the Americas that began in 1492. This suggests that Brazil *L. infantum* population might have been imported during the “Brazilian Gold Rush”, that started in 1690, or by the transfer of the Portuguese court to Brazil in 1808 to flee from Napoleonic troops; both resulting in considerable immigration to Brazil and socioeconomic restructuration of the country. Hence, VL is likely to be one of many diseases introduced to the Americas by Conquistadors and/or Iberian settlers, including smallpox, influenza, typhus, measles, plague, yellow fever and malaria ^21,22^. The large increase in *L. infantum* population size after 1950 in Brazil supports that *L. infantum* dispersal and population expansion occurred recently (Figure 2 C), with the Brazilian urbanisation in the 1960s^23^. Our analysis suggests that Brazil now has the largest effective population size (Ne) and the most rapid expansion of all the endemic regions (Figure 2 C). This is in accordance with the high prevalence of canine leishmaniasis (dogs are the main *L. infantum* reservoir in Brazil) in endemic areas, where from 3% to 36% of dogs are found to be actively infected^24^.

To support the analysis of kDNA phylogeny with BEAST, we conducted a coalescent population size and population split dates estimation using SMC++ ^13^, based on 496,704 segregating sites in the nuclear genome. As SMC++ generation estimates are produced in coalescent units rather than calendar years, we used the kDNA estimations for South-America-Europe samples coalescence date (calendar date: 1867; generations: 4,896) and South Asia and *Leishmania infantum* coalescence date (South and America-European) (calendar date: 872; generations: 40,268) to calibrate SMC++ generations to calendar years (see methods), which resulted in a mean value of 33 generations/year. This estimate of generations/year is in agreement with the rise in population size of the main population of *L. donovani* in the Indian subcontinent in 1850, that was estimated using other methods^9^ (Supplementary figure 5). With this calibration, we estimate the LdSC was present in the East African population as far back as ~12,680 BCE, in close proximity to the start of the Holocene ~11,000 years ago, after the last glacial period. The Holocene was a period with warmer climate, and a rapid proliferation and spread of humans worldwide, with the transition from hunter-gatherer nomads to more sedentary and agricultural lifestyles ^25–27 28^. This demographic expansion and more sedentary lifestyle may have been important for the parasite dispersal and the adaptation of its vector to domestic environments. This is also in accordance with the discovery of *L. donovani* DNA in Egyptian mummies from 2050–1650 BCE, and Nubian human remains in Sudan from 550-1500 CE ^29^.

Our TMRCA estimate is far more recent than previous estimates of one million or ten million years ago that used smaller or more restricted data sets ^30,31^. One source of the discrepancy is the calibration of both previous studies, which assumed a Trypanosomatid divergence of 100 million years ago, coincident with continental divergence ^32^. Current evidence supports a more recent divergence of Trypanosomatid species via the migration of bats ^33–36^. We propose that a more recent dispersal of the LdSC within human history is more consistent than the previous estimates predating the migration of *Homo sapiens* out of Africa 200-300,00 years ago ^37–39^. In this scenario, humans and domestic/farm animals would have been the main drivers of LdSC dispersal.

In summary, our analysis indicates that even though LdSC was already in Africa for at least 12 thousand years, its global dispersal occurred within the last two millennia. The timing of the initial dispersal from Africa (~300 CE) and the seeding of Brazil (~1800 CE) were coincident with population movements facilitated by the Aksumite, Roman, Islamic and Portuguese and Spanish empires. While *L. infantum* appears to have been introduced to South America during times of considerable migration from European colonisation, the population expansion only occurred with the urbanisation of Brazil, since the 1960s.

## Acknowledgements

DCJ was supported by a Wellcome Trust Seed Award in Science to DCJ (208965/Z/17/Z). CAG, JRLC, DCJ were supported by the MRC Newton as a component of the UK: Brazil Joint Centre Partnership in Leishmaniasis (MR/S019472/1) and a MRC New Investigator Research Grant to D.C.J. (MR/T016019/1). CML was supported by the MRC-funded DiMeN PhD program. This project utilised the Viking Research computing cluster at the University of York. We acknowledge the support of The Genomics Laboratory in the Technology Facility, Department of Biology, University of York.

DCJ was supported by a Wellcome Trust Seed Award

## Methods

### Strain culture and genome sequencing

This work was based on newly generated LdSC sequences as well as LdSC sequences from public repositories.

Parasites from the Coleção de Leishmania from Oswaldo Cruz Foundations (CLIOC) were obtained from human and dog patients as part of normal diagnosis and treatment with no necessary invasive procedures and with written and/or verbal consent recorded at the time of clinical examination. The information received and registered by CLIOC guarantees the patients’ anonymity. The access to genetic heritage followed the Brazilian Law of Biodiversity and it was registered at Sisgen (Access number A5C0A32). All strains were cultured in biphasic (Novy–MacNeal–Nicolle [NNN] + Schneider’s) medium prior to DNA extraction. Parasites from The Montpellier Leishmania Collection were cultured in SDM-79 medium supplemented with 10% heat-inactivated foetal calf serum at 25°C for 4 to 7 days. Four isolates were donated by Carlos Robello from Institut Pasteur de Montevideo, Uruguay and cultured as reported in ^40^.

DNA extractions were performed using DNeasy Blood and Tissue kits (Qiagen) according to the manufacturer’s instructions. Genome sequencing was performed on Illumina HiSeq 2500 machines (or similar) to produce paired end 150 nt reads. The samples were sequenced to provide mapped read coverage of ≥20x. Raw sequencing reads were submitted to National Center for Biotechnology Information (NCBI)’s sequencing read archive (SRA) under the BioProject accession PRJNA865107. Sample IDs can be seen on Supplementary Table 1.

Publicly available WGS LdSC data were obtained from NCBI SRA (https://www.ncbi.nlm.nih.gov/sra/). A full list of strain names/SRA numbers can be found in Supplementary Table 1.

### Sequence analysis/variant calling

To remove low quality reads and adapter sequences, WGS reads from each LdSC Isolate were trimmed using fastp v 0.23.2 ^41^, removing reads with quality below Phred 20, shorter than 30 nt and removing bases in the N and C terminal with quality below Phred 25 (--average_qual 20 --length_required 30 --cut_front --cut_front_window_size 1 --cut_front_mean_quality 25 --cut_tail --cut_tail_window_size 1 --cut_tail_mean_quality 25).

The *L. infantum* reference genome (strain JPCM5) was downloaded from TriTrypDB (version 66) ^42,43^. The *L. infantum* maxicircle sequence was downloaded from NCBI (BK010877.1), and combined with the nuclear genome data prior to read mapping. Each polished read library was mapped to the *L. infantum* JPCM5 nuclear genome combined with the Maxicircle (BK010877.1) reference using BWA mem v.0.7.17 ^44^. The mapped reads were filtered by mapping quality 30 using samtools v.1.17 ^45^. Mate-pair data was corrected with samtools fixmate; and PCR duplicates were removed with samtools markdup.

For each isolate, SNPs and indels were called using Freebayes v.1.3.6 (https://github.com/ekg/freebayes), accepting calls with a minimum alternative allele read count ≥5; and octopus v.0.7.2 ^46^. We accepted calls discovered by both methods, merged all single-isolate VCFs and regenotyped with Freebayes. The multi-sample VCF was filtered to contain only biallelic sites, with SNP quality > 30, Minor Allele Count (MAC) of at least 1 (remove non-variant sites), read depth higher than ¼ and lower than 4x the mean coverage of all SNP sites; and mapping quality 30 on both reference and alternate alleles, using BCFtools v.1.15.1 ^47^ (bcftools view -m 2 -M 2 -i ‘QUAL > 30 & MAC >=1 & INFO/DP > 10699 & INFO/DP < 171190 & INFO/MQM >30 & INFO/MQMR >30). Next, we removed Insertion and deletions sites; isolates with more than 5% of missing data and SNP positions with more than 10% of missing data, using VCFtools v.0.1.16 ^48^. The SNP data from the Nuclear genome was separated from the kDNA data using VCFtools v.0.1.16, resulting in a total of 899 isolates, with 496,704 SNP segregating sites (~15 Segregating sites/Kb).

As there was a large variation in the kDNA coverages, the filtering was similar to the one used in the nuclear genome, but without removing SNPs based on coverage, and removing SNPs in the kDNA repetitive region (kDNA positions 17036-18277), using BCFtools v1.15.1 ^47^ (bcftools view -m 2 -M 2 --regions BK010877.1:1-17035 --types snps -i ‘QUAL > 30 & MAC >=1 & INFO/MQM >30 & INFO/MQMR >30’). After removal of isolates with more than 5% of missing data and SNP positions with more than 10% of missing data, a total 874 isolates and 382 SNP segregating sites were obtained for the kDNA data (~22.4 Segregating sites/Kb).

### Population and diversity analysis

To generate the LdSC nuclear genome phylogeny network, the multisample VCF was converted to nexus format using vcf2phylip (https://github.com/edgardomortiz/vcf2phylip), which was used as input for the NeighborNet method in SplitsTree V4.19.2 ^49^. The phylogeny network plot was generated using ggtree ^50^, where each isolate was coloured based on its main admixture population (see below). The same phylogeny network, coloured by the isolation region instead of admixture population can be seen in Supplementary Figure 1.

Population structure (admixture) and PCA analyses used unlinked SNPs (*r*^2^ > 0.5), filtered using PLINK v.2 ^51^ with the options --indep-pairwise 10kb 1 0.5, resulting in a final number of 312,361 segregating sites. Population structure analysis was performed with Admixture v1.3 ^52^, which was run unsupervised, with *K*=1-20 and 10 repetitions. A value of K=9 was selected, based on Cross Validation (CV) analysis. Plots for other values of K can be seen in Supplementary Figure 2. Principal component analysis (PCA) coordinates were produced with PLINK. Both Admixture and PCA plots were generated in R, using ggplot ^53^.

### kDNA phylogeny and dating

Individual kDNA fasta sequences from the 874 isolates incorporating their SNPs were extracted from the VCF using bcftools consensus ^47^, and aligned using MAFFT.v7.453 ^54^. To assess the most likely LdSC root position, we generated a kDNA phylogeny including the sequence from three outgroup samples: *Leishmania major* (LR697138.1), *Leishmania adleri* (LR697136.1) and *Leishmania braziliensis* (LR697134.1), with iqtree2.v2.2.0 ^55^, with the GTR nucleotide substitution model and 1000 SH-aLRT replicates. The tree image was generated in ITOL ^56^, confirming that the root should be in the branch leading to East Africa samples (Supplementary figure 4).

From the 874 LdSC isolates, 567 had recorded dates of isolation, and were used in the dating analysis. Initially the multiple sequence alignment of the fasta sequence of these kDNA sequences was converted to nexus format, using goalign v.0.3.7 ^57^, which was used in BEAST v.10.4 ^12^, with the dated tips method, and different parameters, as seen below. First, to generate the dating of all populations coalescent time, we used the 567 isolates with dated tips, the Bayesian skyline coalescent prior, 8 population size estimates, GTR nucleotide substitution model, fixed clock rate and 500M MCMC iterations, starting from a random tree. This resulted in a rooted tree in East Africa, as expected. The molecular clock was estimated as 2.123×10^−6^/year. Assuming ~ 33 generations/year, we obtain 0.64333e-7, which is in the range of the expected values in eukaryote mitochondria (0.0676 to 1.6 ×10^−7^ changes per site per generation) 58.

Then, we used key datasets to further and independently validate some of the coalescent times, using BEAST: 212 isolates from South America-Europe dataset (206 from Brazil, 10 From Spain, 8 from France, 6 from Italy, 5 from Uruguai, 3 from Portugal, 3 from

Honduras, 1 from Yuguslavia, 1 from Paraguai, 1 from Panama, 1 from Greece, 1 from Albania), using exponential growth coalescent prior, GTR, fixed clock, 100M MCMC replicates, resulting in a root date of 1867 CE; and the same 212 isolates from South America-Europe dataset and 243 isolates from South Asia (123 from Nepal, 109 from India and 11 from Bangladesh), using bayesian skyline coalescent prior, 3 population size estimates, GTR, fixed clock, 100M MCMC replicates, resulting in a mean root date of 872 CE.

### Nuclear population size estimates across time

Past population size changes and population divergence times (both measured in generations) were estimated with SMC++ v1.15.4 ^13^, using populations of isolates from the Admixture analysis: EA1, EA2, SA1, SA2 and Brazil (population lists on Supplementary Table 1). Estimates of population size changes were made with SMC++, with the parameters -- missing-cutoff 500000, --thinning 100, 200 or 1000, mutation rate of 1.99 x 10^−9^ as reported in ^6^, with 5 distinguished lineages, and up to 100 other lineages (depending on the population; Brazil n=100, EA1 n=100, EA2 n=8, SA1 n=12, SA2 n=100). For each population, SMC++ estimate runs were repeated ten times on each subpopulation, specifying distinguished and other lineages selected at random for each replicate. Only generation numbers that were supported by at least 5 of the 10 replicates were evaluated. To convert the generations to dates, we combined the SMC++ generations and the previously assessed kDNA root date for South-America-Europe (root date: 1867; Generations: 4896) and South Asia (root date: 872; Generations: 40268), resulting in a mean value of 33.6 generations/year. To further test this number of generations/year, we estimated the population size estimates for the ISC5 populations described in Imamura ^9^, using smc++, with 5 distinguished lineages, 30 other lineages, and 200 thinning; or 5 distinguished lineages, 30 other lineages, and 1000 thinning; both with 10 replicates. Using 33.6 generations/years as a calibration, we could identify the main ISC5 population expansion in ~1850.

