## Supplementary Figure for "The global dispersal of visceral leishmaniasis occurred within human history"

### Supplementary Figures

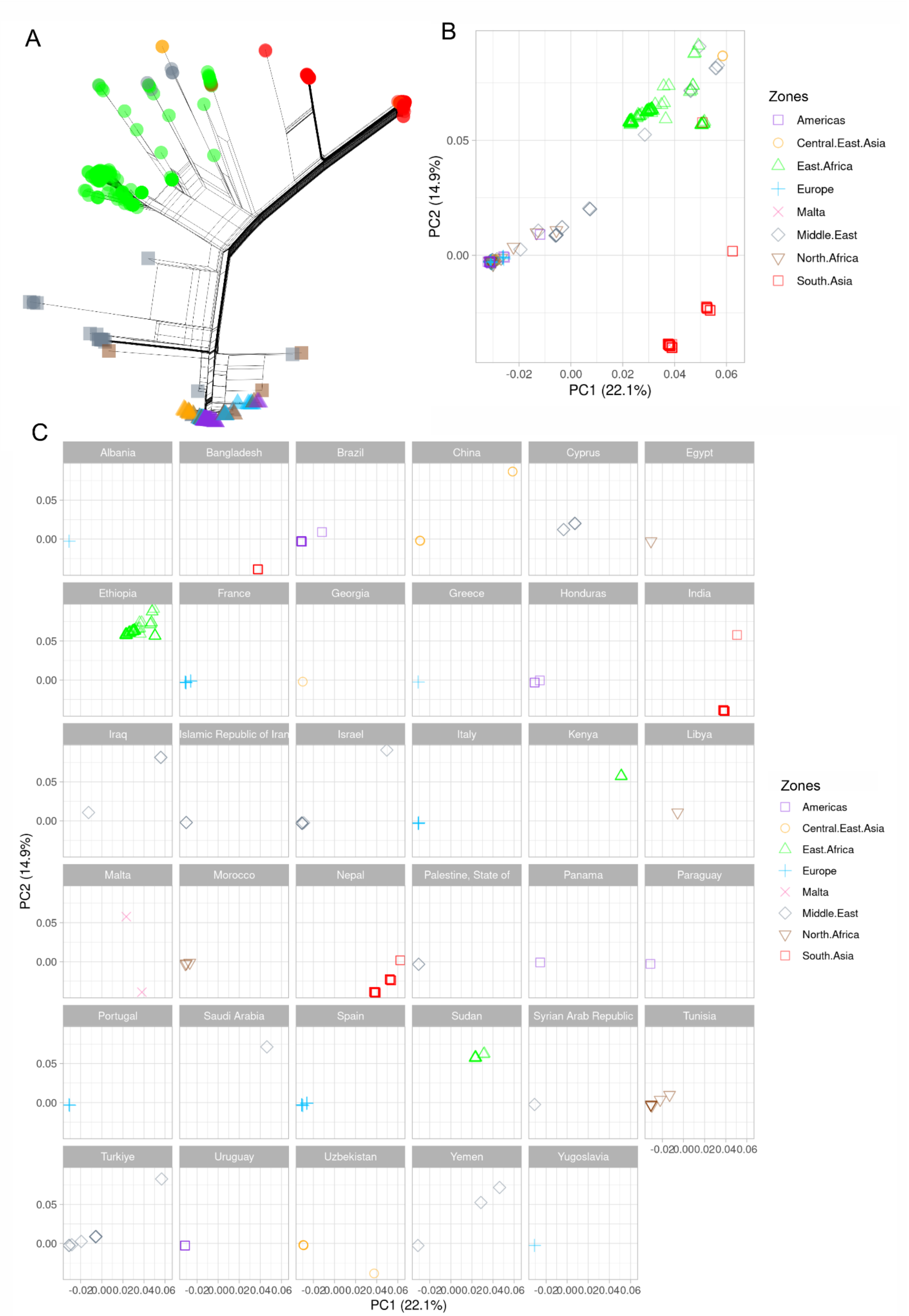


**Supplementary Figure 1: LdSC phylogeny network and PCA coloured by reported local of isolation. A)** Phylogenetic network based on the whole genome sequence of 899 **LdSC**

isolates from 35 countries. Similar to Figure 1 A, but here each isolate is coloured by its zone of origin: Purple: Americas; Yellow: Central.East.Asia; Green: East Africa; Blue: Europe; Pink: Malta; Gray: Middle.East; Brown: North.Africa; Red: South.Asia. **B)** PCA of the 899 LdSC coloured by Zones. **C)** PCA of the 899 LdSC isolates coloured by zone. Each country of isolation is represented in a different panel, with the same X and Y coordinates as in **B**.

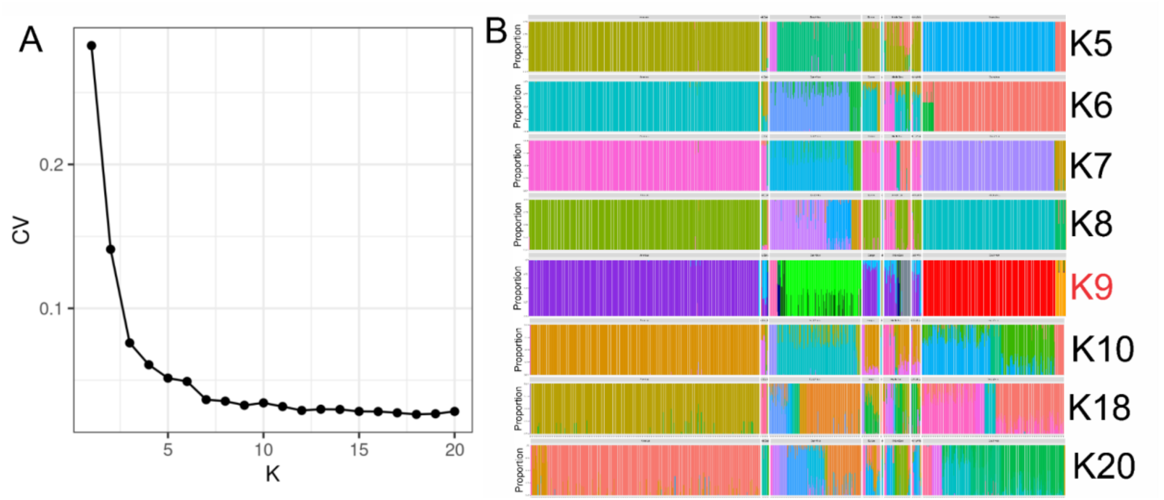


**Supplementary Figure 2: Nuclear genome population structure results. A)** Admixture Cross Validation Error (CV) for the different K values. **B)** Admixture plot for the 899 LdSC isolates using different K values. The K9 is highlighted in red.

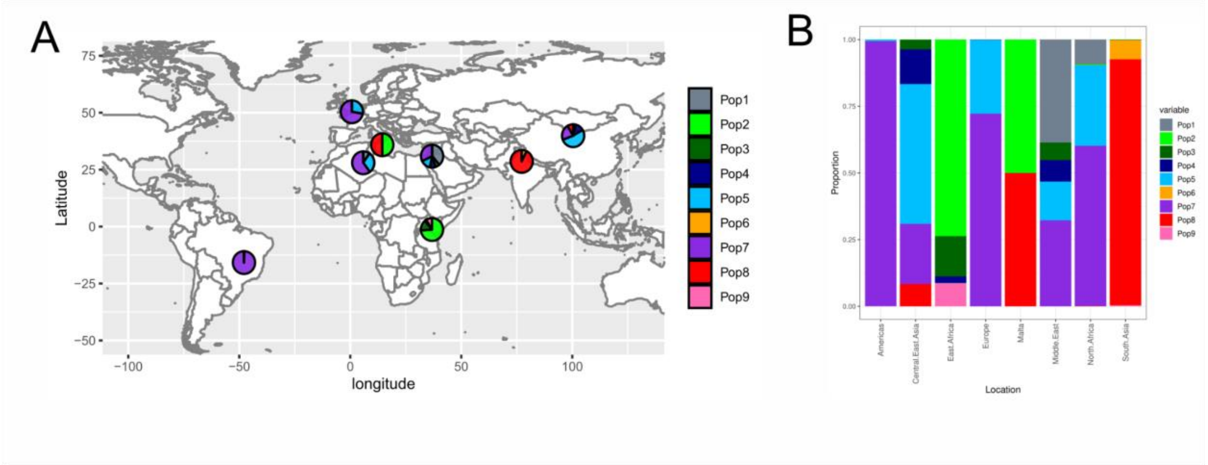


**Supplementary figure 3: Geographic distribution of all evaluated LdSC isolates. A)** Map representing the proportion of isolates from each population in each geographic region. **B)** Admixture plot showing the proportion of each population in each geographical zone.

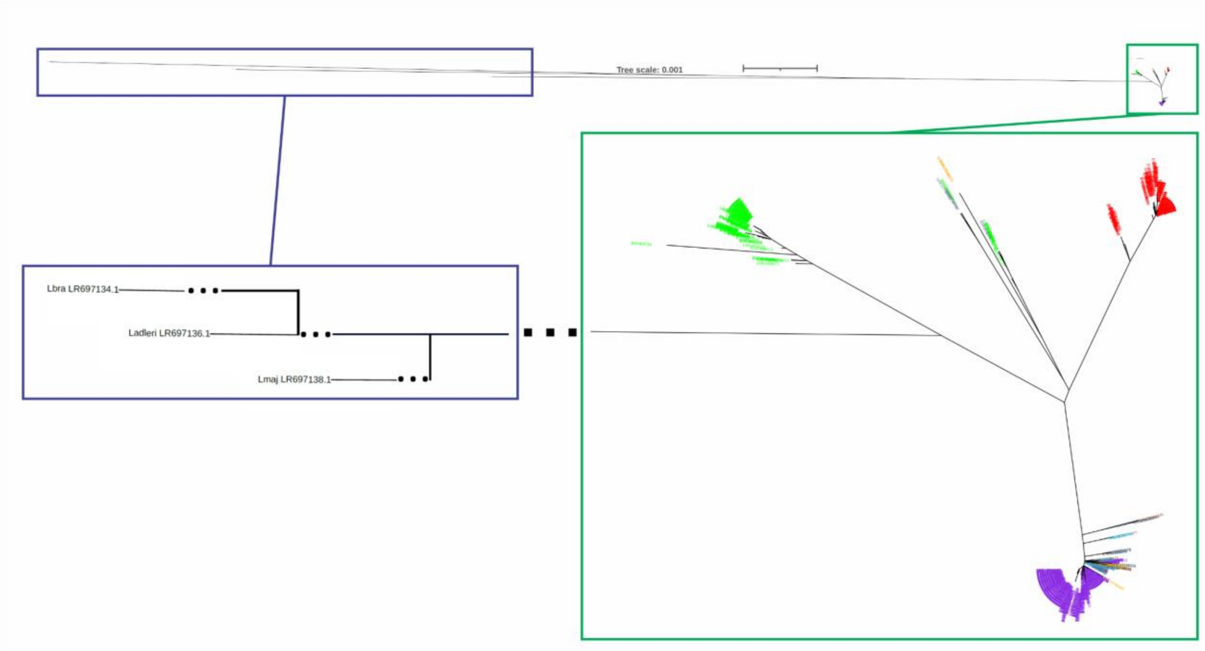


**Supplementary figure 4: kDNA phylogeny with three outgroups.** Maximum likelihood phylogeny based on kDNA SNPs, including three outgroup sequences: *Leishmania major* (LR697138.1), *Leishmania adleri* (LR697136.1) and *Leishmania braziliensis* (LR697134.1), using iqtree2, with the GTR nucleotide substitution model and 1000 SH-aLRT replicates. The top panel corresponds to the phylogram, showing all evaluated distances. The botton panel zooms in the main three (green box) and outgroups (purple box) showing that the root should be included within the East Africa branch. This is in accordance with the BEAST root estimation in Figure 2. The isolates are coloured based on the reported location of isolation.

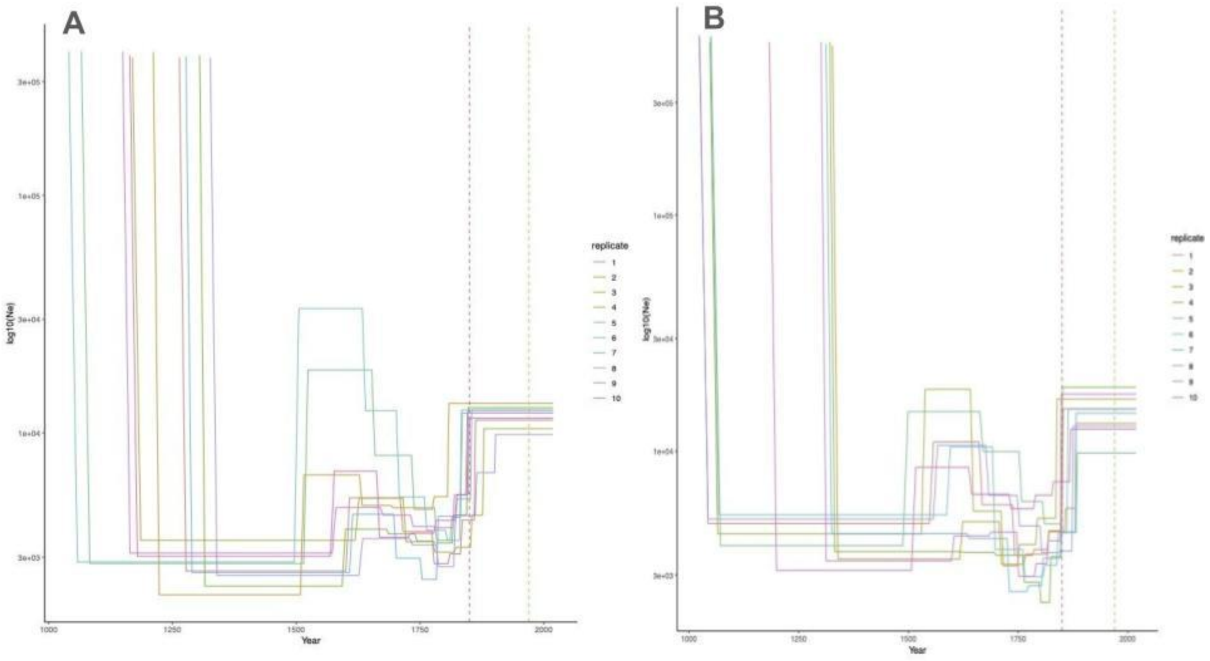


**Supplementary figure 5: SMC++ population size estimates for the main population of *L. donovani* in the Indian subcontinent.** Using samples from the ISC5 population as defined by Imamura (Imamura et al. 2016), smc++ (Terhorst et al. 2017) was run with A) 5 distinguished lineages, 30 other lineages, and 200 thinning and B) 5 distinguished lineages, 30 other lineages, and 1000 thinning. The coloured lines represent 10 independent replicates each with 5 distinguished lineages and 30 other lineages chosen at random from the samples. The red dotted line represents the year 1850, the approximate date of the main population expansion from Imamura *et. al.* The green dotted line represents the year 1970, the approximate date of the ISC5 population expansion estimated in Imamura *et al*. These SMC runs do not detect the 1970 population expansion. However, using our estimate of 33.6 generations/year does detect the rise of the main population in 1850.
